# Flavor-Cyber-Agriculture: Optimization of plant metabolites in an open-source control environment through surrogate modeling

**DOI:** 10.1101/424226

**Authors:** Arielle J. Johnson, Elliot Meyerson, John de la Parra, Timothy L. Savas, Risto Miikkulainen, Caleb B. Harper

**Affiliations:** Media Lab, Massachusetts Institute of Technology, Cambridge, Massachusetts, United States of America; Department of Computer Science, University of Texas at Austin, Austin, Texas, United States of America; Cognizant Technology Solutions, San Francisco, California, United States of America; Harvard University Herbaria, Harvard University, Cambridge, Massachusetts, United States of America

## Abstract

Food production in conventional agriculture faces numerous challenges such as reducing waste, meeting demand, maintaining flavor, and providing nutrition. Contained environments under artificial climate control, or cyber-agriculture, could in principle be used to meet many of these challenges. Through such environments, phenotypic expression of the plant---mass, edible yield, flavor, and nutrients---can be actuated through a “climate recipe,” where light, water, nutrients, temperature, and other climate and ecological variables are optimized to achieve a desired result. This paper describes a method for doing this optimization for the desired result of flavor by combining cyber-agriculture, metabolomic phenotype (chemotype) measurements, and machine learning. In a pilot experiment, (1) environmental conditions, i.e. photoperiod and ultraviolet (UV) light (known to affect production of flavor-active molecules in edible plants) were applied under different regimes to basil plants (*Ocimum basilicum*) growing inside a hydroponic farm with an open-source design; (2) flavor-active volatile molecules were measured in each plant using gas chromatography-mass spectrometry (GC-MS); and (3) symbolic regression was used to construct a surrogate model of this chemistry from the input environmental variables, and this model was used to discover new combinations of photoperiod and UV light to increase this chemistry. These new combinations, or climate recipes, were then implemented in the hydroponic farm, and several of them resulted in a marked increase in volatiles over control. The process also led to two important insights: it demonstrated a “dilution effect”, i.e. a negative correlation between weight and desirable chemical species, and it discovered the surprising effect that a 24-hour photoperiod of photosynthetic-active radiation, the equivalent of all-day light, induces the most flavor molecule production in basil. In this manner, surrogate optimization through machine learning can be used to discover effective recipes for cyber-agriculture that would be difficult and time-consuming to find using hand-designed experiments.

## 1 Introduction

The so-called “dilution effect,” noted since the 1940’s and systematically reviewed since the early 1980’s [1], describes an inverse relationship between yield and nutrient concentration in food: For many nutritionally-important chemical constituents of food plants, such as minerals, protein, and vitamins, an increase in biomass is accompanied by a decrease in nutrient concentration. This effect has been systematically demonstrated in historical nutrient content studies over the last 50-70 years [2,3], as well as in controlled side-by-side trials that have shown a relationship between nutrient dilution and genetics [4], artificial fertilization [5], and elevated carbon dioxide levels related to climate change [6,7]. Flavor, known to be an important element of food and of eating behavior for organisms from insects to humans [8], has been declining alongside nutrients over approximately the last 50 years [9-11] in inverse proportion to rising yields. Declining flavor is of concern because flavor-active molecules in plants frequently have either positive health benefits themselves (antioxidant, antimicrobial, anti-inflammatory) themselves or signal the presence of other beneficial or essential molecules, for example by being the enzymatic products of precursors necessary for human health and nutrition (e.g. pro-vitamin A carotenoids, essential amino or fatty acids) necessary for human nutrition and health [9].

Vertical farming, or more generally cyber-agriculture, is a plant-growing format that employs contained environments where light, water, nutrients, temperature, and other climate variables are provided artificially under computer control [12-14]. Data from environmental sensors is used to actuate climatic conditions according to a “recipe” designed for best possible outcome such as largest yield, best flavor, desired nutrients, and least cost. With cyber-agriculture, in principle it may be possible to increase quality and quantity of food production, minimize waste and cost, and grow food with optimized climate recipes anywhere including locations otherwise unable to support agriculture. Conventional agriculture has been optimized for yield. What if it were optimized for quality and flavor?

This paper describes a proof-of-concept method aimed at optimizing flavor in a cyber-agricultural controlled environment, and a pilot experiment to validate this method. An experimental container, called the Food Computer (FC) [12], was built at the Massachusetts Institute of Technology (MIT) Media Lab with sensors, actuators, and computer control. Basil (*Ocimum basilicum*) was chosen as the model organism because it has a fast growth cycle (five weeks), and because the outcome can be readily measured in terms of fresh weight (quantity), and chemical analysis of flavor (quality). To keep the optimization problem manageable, it focused on the lighting conditions, keeping the other variables constant. A number of known recipes were first implemented, together with a broad range of their variations [15]. Machine learning technology [16-18] was then used to optimize these recipes further: based on these recipes and their associated outcomes, a surrogate model was first constructed using symbolic regression. The surrogate model was then searched to discover potentially better lighting recipes, which were then tested in the experimental container. Indeed, recipes that yielded significantly better flavor were discovered in this process. In addition, the results demonstrated the dilution effect, and a new, surprising positive effect of 24-hour light. The experiments thus demonstrated that cyber-agriculture is a potentially viable solution to several problems that agriculture faces today.

## 2 Methods

In this section the problem of flavor optimization is first defined, the Food Computer environment for controlled growth experiments is then described, and finally the methods for building a surrogate model and discovering improved growth recipes with it is described.

### 2.1 Measuring and optimizing flavor

Flavor is largely a phenomenon of olfaction [19], and many aroma molecules are produced by the specialized metabolism of plants. Plants have a particularly rich specialized metabolism [20], a set of biosynthetic pathways synthesizing molecules that are not essential for the basic processes of life (cell division, reproduction, etc.) but rather confer fitness and adaptive advantage to the organism in its ecological niche [21], related to stress tolerance, defense, and communication [22]. Their expression and induction depend, to various degrees, on environmental and ecological conditions [23].

Cyber-physical agriculture methods such as the Food Computer, where data from environmental sensors informs the actuation of climatic conditions according to a climate recipe [12-14] present unique opportunities for inducing plant phenotypic changes through environmental/ecological conditions alone. One example of this approach is to apply the ecological stresses to which adaptations have evolved as specific biosynthetic pathways.

The basil plant, *O. basilicum*, is typical of herbaceous plants in that it produces many aromatic molecules, particularly the terpenoids 1,8-cineole, linalool, camphor, borneol, bergamotene, and farnesene, and the phenylpropenes eugenol, methyleugenol, and estragole [24]. These molecules are thought to play varying roles in stress adaptation and defense, and the production by the basil plant of aromatic molecules has been shown to increase upon exposure to these stresses, including water stress [25], ultraviolet (UV) and photosynthetic-active radiation (PAR) light [26–28], heat [29], bacteria [30], chitosan (a compound derived from chitin, found in insect exoskeletons and fungal cell walls, [31]), and sodium and other minerals [32].

This paper explores methods for increasing flavor molecule production in *O. basilicum*, using: (1) UV light, PAR, and photoperiod as environmental and stress variables; (2) gas chromatography-mass spectrometry (GC-MS) for semiquantitative analysis of volatiles; (3) surrogate optimization for discovering conditions that will maximize production of these volatiles.

### 2.2 Controlled growth environment

This section describes the design of the Food Computer, i.e. the physical container environment used in the pilot experiment with basil. It also describes the process for growing basil in this environment, and methods for measuring the growth outcome in terms of weight and chemistry.

#### 2.2.1 Food computer

All basil plants were grown in a Food Server (Fig 1), a multi-tray, multi-rack hydroponic configuration of the OpenAg Food Computer™ environment [12]. Basil plants were germinated in engineered foam rooting cubes (Oasis Grower Solutions, Kent, OH), then transplanted to 36-position (4×9) food-grade resin floating lettuce rafts (Beaver Plastics, Acheson, AB, Canada) at 14 days of age. The plants were grown in a shallow water culture hydroponic system according to the details in Table 1.

**Table 1.**
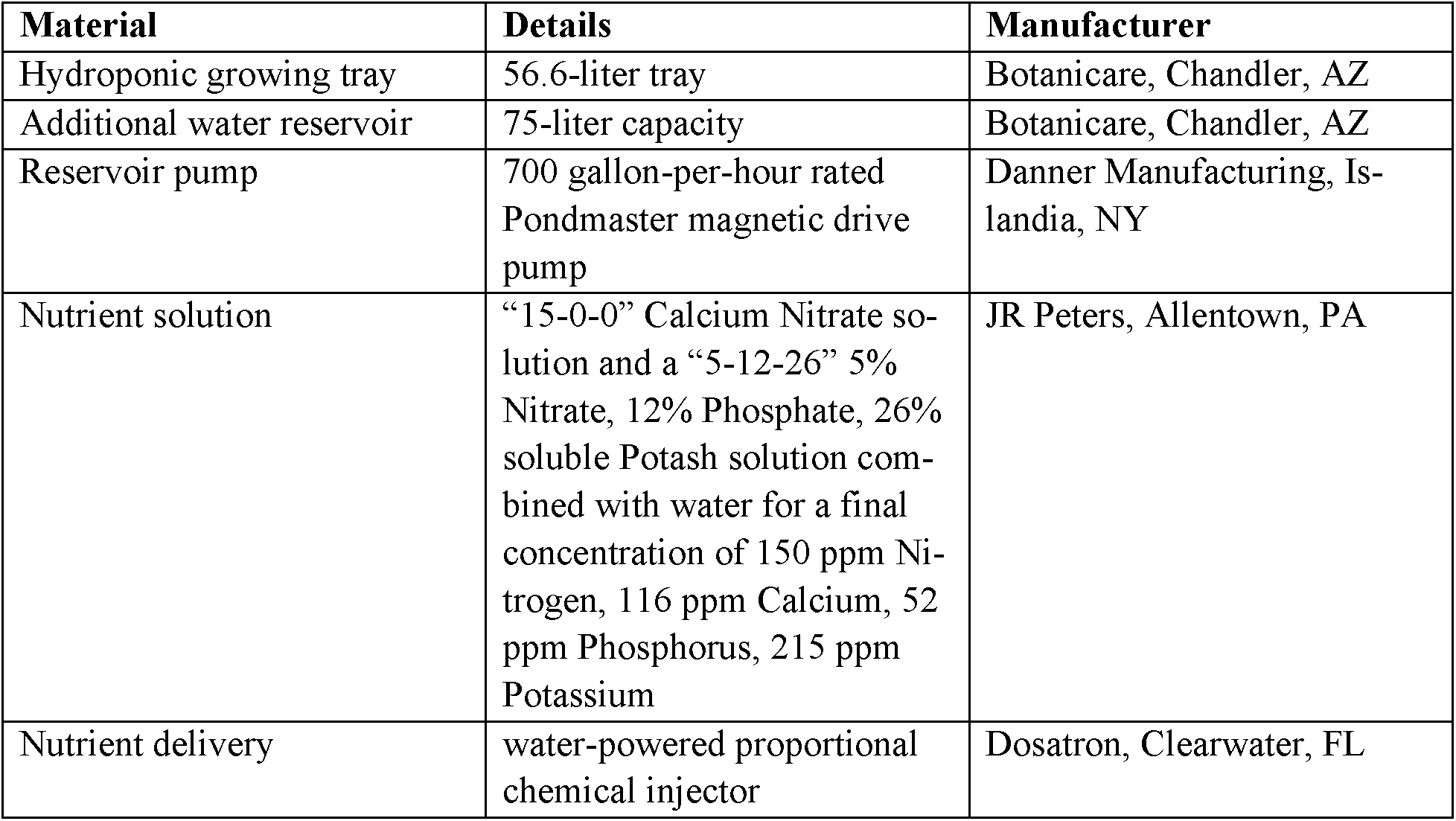
Hydroponic system design elements

**Fig 1.**
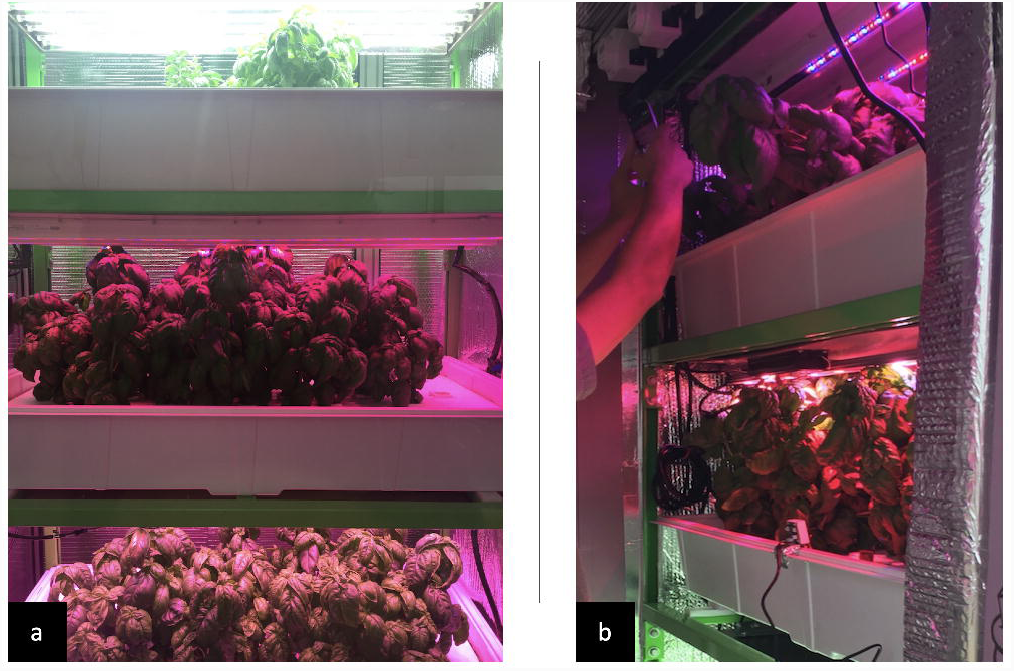
MIT Media Lab Food Server. (a): Growing configuration inside the Food Server. (b): A view inside the Food Server during experimentation.

The Food Server was set up with trays in vertical stacks of three (denoted 0, 1, and 2) within a custom designed unit according to the elements described in Table 2.

**Table 2.**
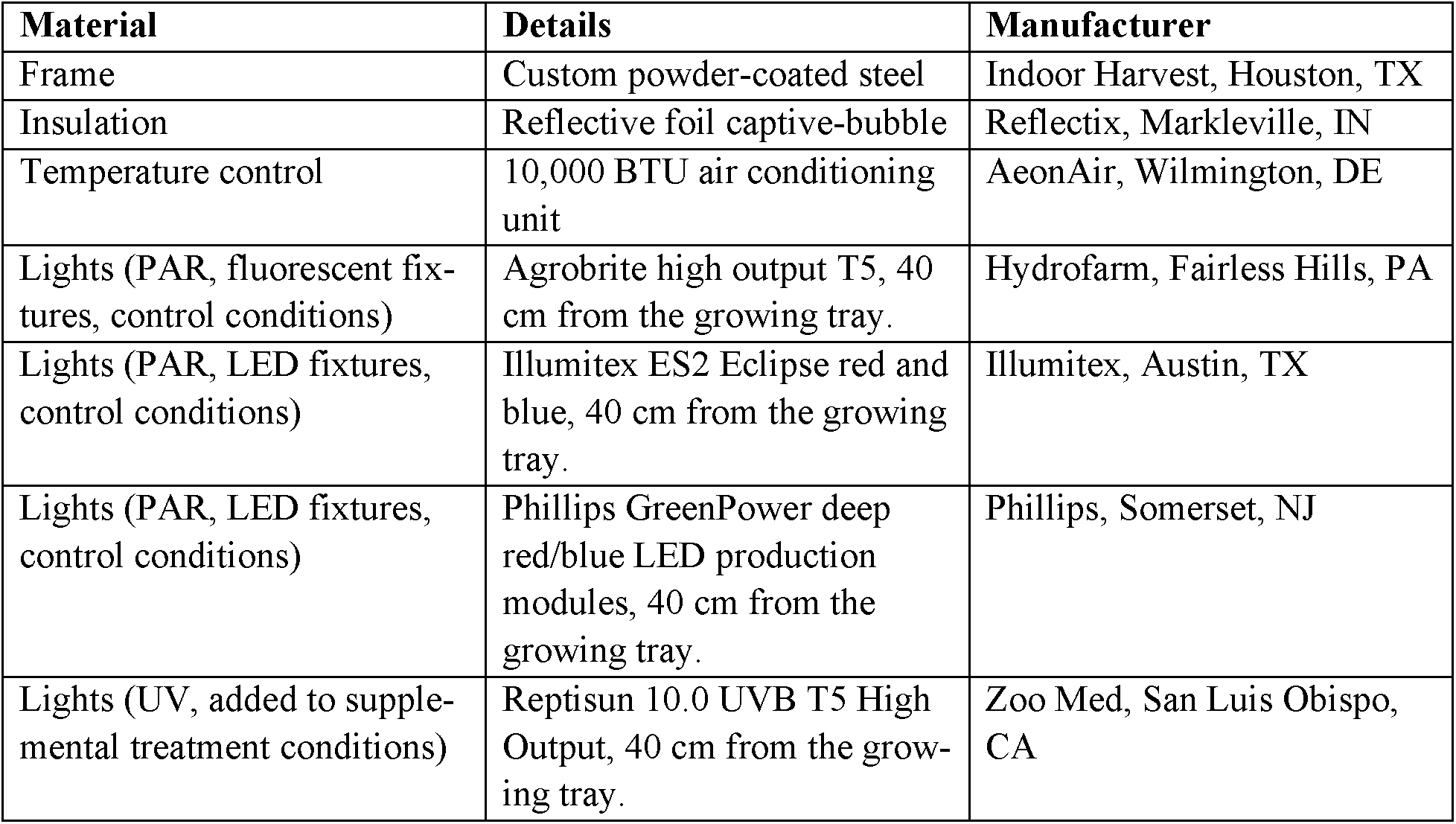
Food Server environmental design elements

#### 2.2.2 Plant species and climate recipes

Common Sweet Basil (*O. basilicum var “*Sweet”) seeds (Eden Brothers, Arden, NC) were used in the pilot experiment. From 14 days of age to harvest, they were grown in identical trays as described in “Food Computer*”* above, with one of three control condition light fixtures as the only source of PAR (Table 2). Control conditions were grown with the PAR light fixture only; experimental treatment conditions had supplemental UV light. Treatment conditions, or “Climate Recipes”, in Rounds 2 and 3 of the experiment were determined based on suggestions from the surrogate optimization of chemscore from the previous round. The data from Round 1 determined the conditions of Round 2, and the data from Round 2 determined the conditions of Round 3).

#### 2.2.3 Harvest, weight, and length measurement

All plants in each round of the experiment were harvested on the same day. Four plants from each treatment condition were used for volatile analysis and the remaining 32 were used for height and weight measurements. Weight measurements were taken with roots removed.

#### 2.2.4 Sampling and sample preparation

Immediately after harvesting, leaves were sampled from four plants from each treatment condition. Fifteen leaves from each plant were harvested: five from near the base, five from the middle, and five from the top, with each set selected randomly. Leaves were immediately frozen with dry ice or liquid nitrogen, homogenized into a powder, and kept frozen. The amount of 1 gram of frozen plant tissue was transferred to a 20 mL amber glass headspace vial (Supelco, Bellefonte, PA) and 2 mL of saturated, cold calcium chloride solution in distilled water was added to prevent enzymatic reactions. The vials were capped with magnetic, polytetrafluoroethylene (PTFE) -lined silicone septa headspace caps (Supelco) and kept on ice before being transferred to the GC-MS.

#### 2.2.5 Volatile analysis

The method of Johnson et al. [33] was adapted for the experiment. Sample vials were placed in the tray of the Gerstel MuliPurpose Sampler 2 (MPS2) autosampler (Gertsel, Linthicum, MD), which performed the extraction and injection. Each vial was individually warmed to 40°C and agitated at 500 rpm for 5 minutes directly before extraction. A conditioned, 2-cm long 50/30 µm-thick polydimethylsiloxane/ divinylbenzene (PDMS/DVB) solid-phase microextraction (SPME) fiber (Supelco) was introduced into the headspace of the vial for 45 minutes at 40°C with rotational shaking at 250 RPM. The fiber was removed from the headspace of the vial and immediately introduced into the inlet of an Agilent model 7890 single quadrupole GC-MS (Agilent Technologies) with a (5%-Phenyl)-methylpolysiloxane (DB-5) column (30 meters long, 0.25 mm internal diameter (i.d.), 0.25 μm film thickness, J&W Scientific, Folsom, CA). The inlet was held at 250°C with a 2:1 split and had a 0.75mm i.d. SPME inlet liner installed (Agilent Technologies). The carrier gas was helium, at a constant flow rate of 1 mL/minute. The starting oven temperature was 40°C, held for 3 minutes, followed by a 2°C/minute ramp until 180°C was reached, then the ramp was increased to 30°C/minute until 250°C was reached, and held for 3 minutes. The total runtime was 47 minutes. The transfer line to the mass spectrometer was held at 250°C, the source temperature was 230°C, and the quadrupole temperature was 150°C. The mass spectrometer had a 1.5-minute solvent delay and was run in scan mode with Electron Impact ionization at 70eV, from *m/z* 40 to *m/z* 300.

Compounds were identified and recorded based on a 90% or higher match using the National Institute of Standards and Technology (NIST) Mass Spectral Database and a signal to noise ratio above 10. Analyte peaks were integrated on the Total Ion Chromatogram (TIC).

#### 2.2.6 Optimization metric: Chemscore

Optimizing the target metric should correspond to maximizing flavor in a general sense. The metric should also be robust to noise, since the number of evaluations is limited, and low-dimensional to make optimization easier.

Basil, like most foods, contains multiple molecules contributing to flavor. An average GC-MS chromatogram of basil contains around 30-40 different identifiable volatile molecules, with concentrations varying over several orders of magnitude. To construct a single metric to optimize, this GC-MS data is aggregated across samples and chemicals as the chemscore. This score is a weighted average of the volatile profile compared to the control condition. It is a holistic placeholder for how flavorful a sample is, while normalizing for varying scales and distributions of different chemicals. Seventeen chemicals common across all GC-MS measurements were selected for the calculation of chemscore.

#### 2.2.7 Comparison metrics: R-Score and Z-Score

For further comparison, an R-Score and a Z-Score, across all volatiles in a sample, were calculated for each treatment condition. The R-Score, the average ratio of volatiles in a treatment condition over their average in the control conditions in a round of the experiment, facilitates comparison of results across the three rounds of the experiment, under the assumption that uncontrollable environmental differences across rounds are captured in differing control results. The Z-Score, which compares the abundances of each volatile molecule in a sample over or below its average in all samples in a round, gives a sense of the overall spread of results in the experiment.

### 2.3 Surrogate optimization

In optimization settings where the target function is expensive to evaluate (either temporally or financially), e.g., in the case of growing basil to maturity, surrogate-based optimization is a common method for minimizing the number of evaluations required to achieve an acceptable solution [34–36]. To choose the next samples to evaluate, surrogate methods build an explicit predictive model of the solution landscape and select the most promising samples according to this “surrogate model’’. To implement such a method, input variables need to be defined, a class of regression models needs to be selected, and a method for discovering the next samples (recipes) from these models needs to be developed. This section details the development of these choices for the experiment in this paper, and notes methods for scaling up future work. A flowchart of the methodology is shown in Fig 2.

**Fig 2.**
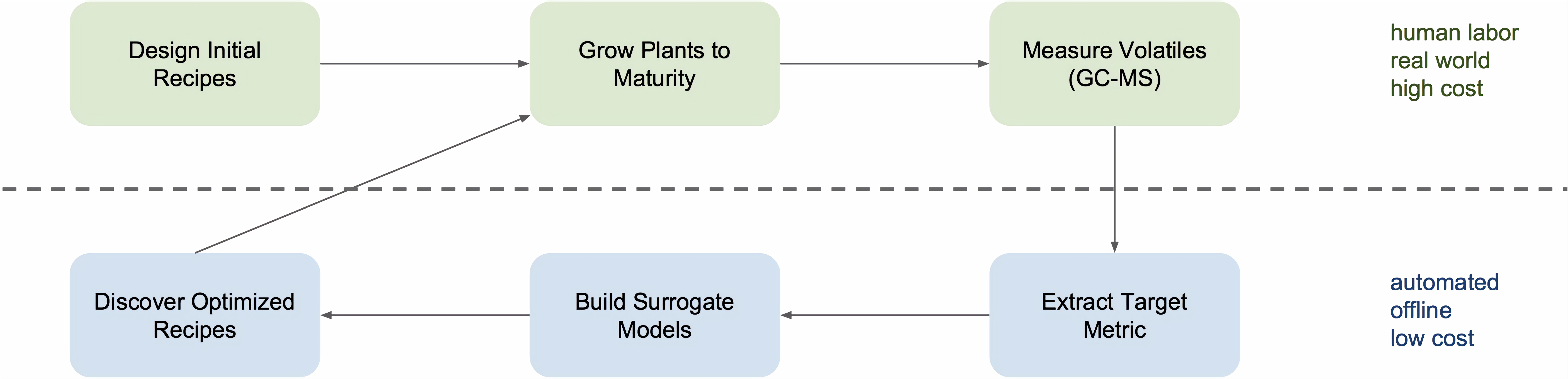
Overview of recipe optimization methodology. First, experimenters Design Initial Recipes based on prior knowledge about the space of acceptable growing conditions. This design includes specifying the input variables and ranges that define the space of possible recipes. Second, these recipes are implemented in real-world controlled environments which Grow Plants to Maturity. Third, GC-MS is used to Measure Volatiles in mature plants. Fourth, this chemical data is aggregated to Extract Target Metric, e.g., chemscore, which is an overall indicator of flavor content. Fifth, the target metric results are used to Build Surrogate Models that model the target metric based on the input recipe variables. Sixth, a search procedure is used to Discover Optimized Recipes that are the most promising for increasing flavor according to the surrogate models. These new recipes are then implemented in the real world as the cycle repeats. The power of this method comes from the fact that modeling and optimization of flavor is done offline with automatically-built models to minimize real-world costs.

#### 2.3.1 Design variables

For this experiment, a recipe was defined by three design variables: photoperiod, UV period, and PAR. Three design variables constituted an appropriate dimensionality for this pilot experiment, following the general rule-of-thumb that, for surrogate-based methods, the number of evaluations required to achieve reasonable results is around ten times the number of dimensions [34]. These variables were chosen because they are already known to increase the accumulation of volatiles [26–28] and are relatively simple to control in the described hardware setup.

Photoperiod is the number of hours the primary light panel is turned on each day. Recipes can thus have photoperiod values anywhere from 0 to 24 hrs. Photoperiod is known to have significant effects on the accumulation of biomass and leaf area in plants [37], as well as the formation of trichomes, the structures that store flavor-active volatiles, in *Thymus vulgaris* (thyme) [38]. *T. vulgaris* and basil are closely related members of the same botanical family, Lamiaceae. In addition, photoperiod has been shown to change the volatile profile of basil [39].

UV period is the number of hours per day plants receive supplemental UV-B radiation. Like photoperiod, UV period can take on values anywhere from 0 to 24 hrs. UV has previously been shown to increase volatile content in basil [26]; it is included so that its effects can be validated and optimized in the Food Computer hardware setting.

PAR is the amount of light available for photosynthesis. In the Food Computer setup, the PAR is determined by the primary light panel. There were nine light panels, each with a unique PAR value. To set PAR values for a batch of nine recipes, one light panel was assigned to each recipe. Thus, in contrast to photoperiod and UV period, each available PAR value can be used only once in each batch. This kind of hardware resource matching constraint is not common in either computer or physical experiments, so a custom optimization method must be developed.

#### 2.3.2 Surrogate model

Symbolic regression [40–42] was used to build surrogate models for predicting a chemscore from the input variables. Symbolic regression uses evolutionary optimization to discover nonlinear algebraic expressions that serve as surrogate models. For the experiment in this paper, a multi-objective Pareto optimization procedure was used [43,44]. The first objective is to minimize error, i.e., mean squared error (MSE) with respect to predicting chemscore; the second objective is to maximize parsimony, i.e., minimize the size of the algebraic expression (number of nodes). The fitting procedure then yields a Pareto front of models, from which a new batch of recipes can be selected.

For the flavor-optimization problem, symbolic regression has several advantages over other popular choices for surrogate models. First, by optimizing for error and parsimony simultaneously, the search is biased towards the kinds of compact algebraic expressions that are desirable in the natural sciences [44]. These expressions are more interpretable than other regression models because the relationships between variables can be read off directly from the expression. Such interpretability can lead to a better understanding of the search space, which helps in developing better models for future experiments.

Second, whereas surrogate models such as Gaussian processes can only interpolate, symbolic regression can extrapolate. Interpolation is sufficient when iterative incremental improvement can eventually lead to an optimal solution. However, in the experiment in this paper, only a single parallel batch of recipes is selected via surrogate optimization to be implemented in the Food Computer. So, it is advantageous to consider strong optimistic predictions a model makes about sparse regions in the recipe space. Note that if this process were used over multiple iterations, an inordinate amount of resources could be spent at the extremes of the recipe space.

Third, symbolic regression is robust to normalization of input and output variables: It automatically discovers reasonable scaling factors to use through optimized constants that are found to be useful in model expressions.

It is important to note that symbolic regression can have significant drawbacks as well [43]. First, it is computationally expensive compared to other regression methods; however, in this paper, computation time is negligible compared to the time it takes to grow a batch of basil recipes. Second, surrogate optimization with symbolic regression models currently lacks theoretical convergence guarantees and performance bounds. Such convergence guarantees have potential practical benefits over many iterations of surrogate optimization; however, since only a single such iteration is performed in the experiment in this paper, such guarantees are unnecessary.

#### 2.3.3 Optimization process

There were three rounds of growing experiments. In each round, there are nine trays of basil growing in parallel. To ensure consistency across rounds, three of these nine trays are fixed to control recipes. This setup leaves six non-control recipes to be selected.

In the first round, recipes were selected by hand [15] to investigate the effects of UV supplement and choice of light panel. To add the photoperiod dimension, and create initial diversity in the recipe space, recipes in the second round were chosen by an unsupervised method: Six non-control recipes were found as centroids of Voronoi tessellation (CVT) given the first round of recipes [45]. Following a trust region approach [35], to implement the bias that good solutions are likely to be relatively close to expert hand-designed recipes, values for each dimension were constrained to be with a constant distance of previously evaluated values.

In the third round, recipes were selected from symbolic regression surrogate models [46]. Each run of symbolic regression yields a collection of models on the error-parsimony Pareto front. These models were clustered to determine an error threshold above which models were underfitting. The six most parsimonious models not underfitting were then used to define a recipe to run in parallel. Since the recipe space has only three dimensions it is computationally efficient to use a dense grid search to select a recipe that maximizes expected chemscore. Greedy sequential selection is the most popular approach to constructing parallel batches from surrogates [47,48]. The recipes were thus selected sequentially in increasing order of model error. Such a selection handles the constraint that each available PAR value can be selected only once per round. If a variable is ignored by a model, the value of the variable is set to maximize exploration, since the model has indicated that exploitation of this variable is currently not useful.

In the model-building step, symbolic regression was run for 1000 generations, with 2000 models evaluated per generation. Therefore, two million symbolic regression models were evaluated. To find optimal recipes for each of the resulting surrogate models, the surrogate was evaluated for each point in a dense grid with a side length of 100; thus each approximate model was evaluated one million times. The eighteen most promising recipes discovered in this surrogate optimization process were then evaluated in the real-world growing experiments.

## 3 Results

The experimental conditions as well as the resultant average weights, R-Scores, chemscores, and Z-Scores are presented in Table 3. The Round 3 rows of Table 3 include additional R-Scores with imputed data. This is because data for one control condition in Round 3 (the last row in Table 3) was lost in the experiment. Imputed values for each chemical for the missing control treatment in Round 3 were computed by regression, i.e., by solving a fully-determined linear system that predicts the value of the third control from the other two, based on the values of the controls in the previous two rounds. Assuming control results are consistent within each round, these additional values make the results easier to compare across rounds.

**Table 3.**
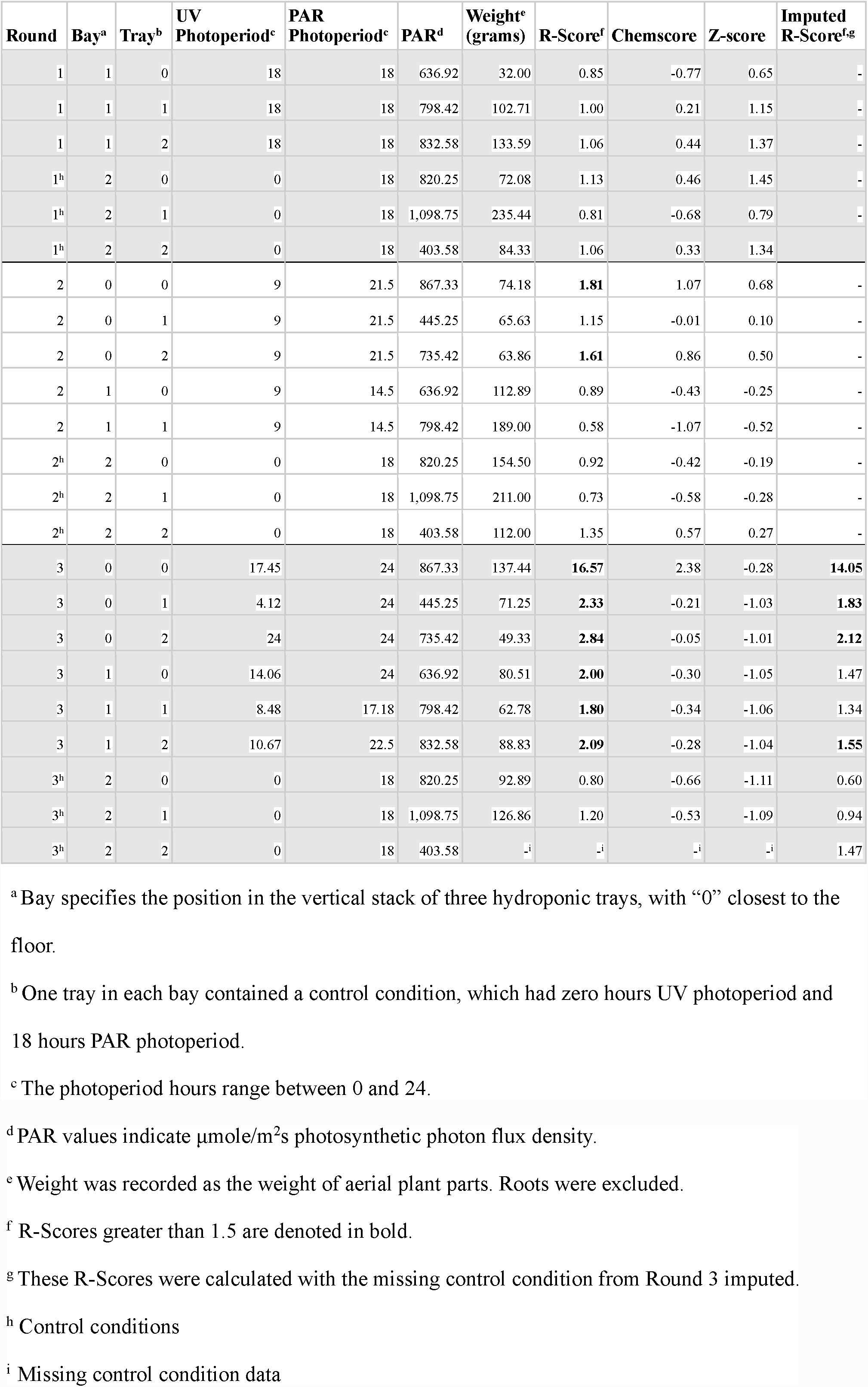
Treatment conditions (UV and PAR photoperiod), weight, and chemical results.

Table 4 gives the correlations between input variables and metrics (Spearman, to account for nonlinearity in the metrics). All of the metrics (R-Score, Weight, Chemscore, Z-Score) are monotonic functions for which a larger number is favorable. Since there is no prior expectation that these metrics have linear scales, the Spearman correlation is used instead of the Pearson correlation. Correlations larger than 0.45 are in bold to show a qualitative separation, as these are above the critical value for a Spearman correlation with 18 samples and p<0.05. Note in particular that the R-Scores are negatively correlated with weight: Optimizing for flavor thus results in smaller plants, and larger plants have less flavor, thus illustrating the “Dilution effect.”

**Table 4:**
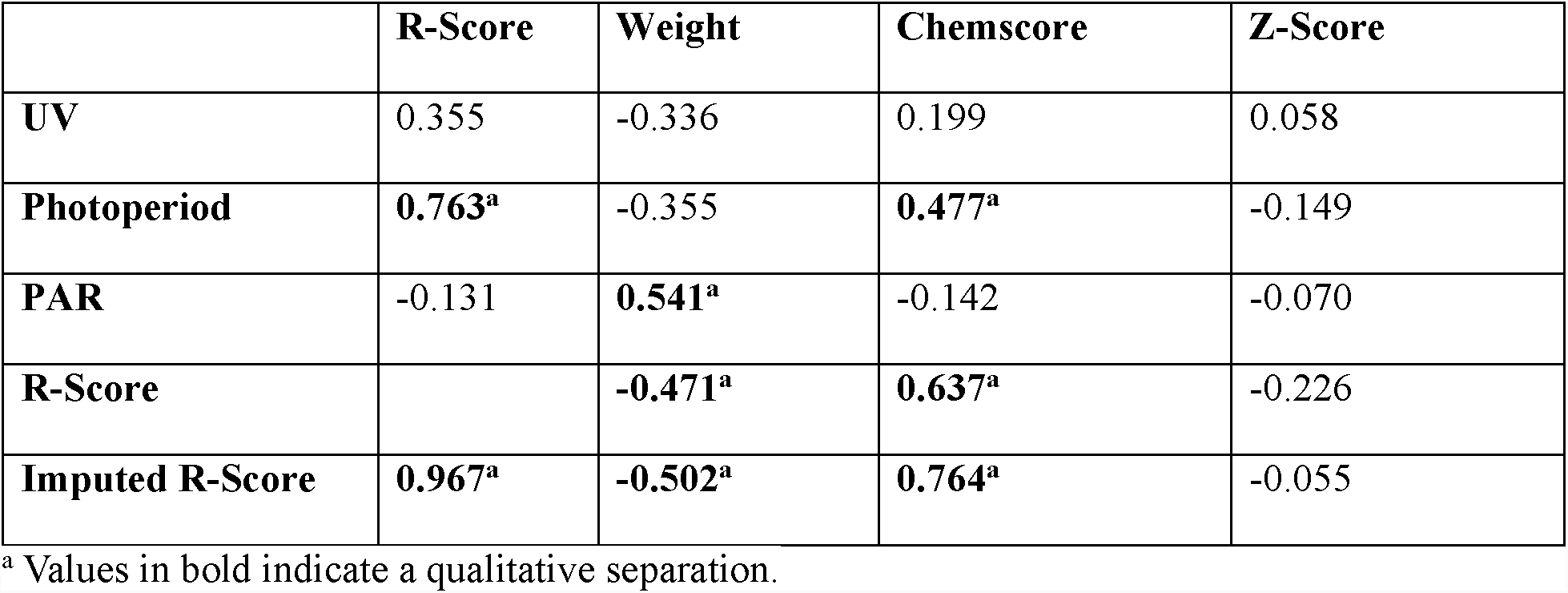
Spearman correlations between selected input variables and metrics.

In the first round, where an 18-hour PAR photoperiod and an 18-hour UV photoperiod were selected by hand, R-Score and chemscore did indicate that UV light or photoperiod increases volatiles. In the second round, two R-Scores (both with UV light and extended PAR photoperiod of 21 hours) were above 1.5, meaning that volatiles holistically increased 50% over control. In the third round, several conditions resulted in an R-Score that met or exceeded this threshold, with many conditions (all with PAR photoperiods of 22.5-24 hours and UV periods of 4-17 hours) doubling the volatile profile compared to control. The discovery of the recipes in Round 3 from the model is illustrated in Fig 3.

**Fig 3.**
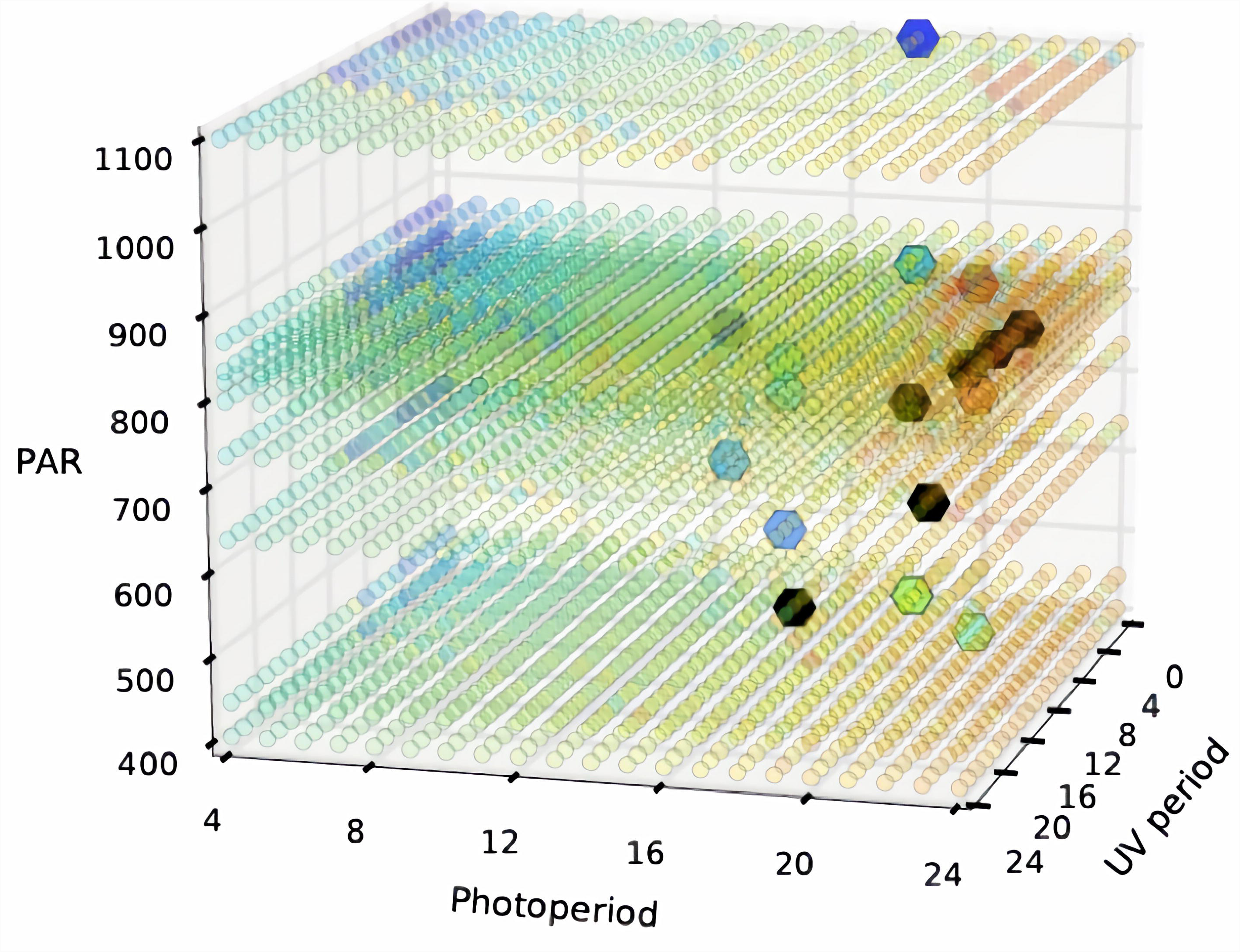
An illustration of the surrogate model and the recipes suggested by the optimization. The three axes correspond to the three actuators and the color of the small dots indicates their value predicted by the model (i.e. flavor; red > yellow > green > blue). The large dots are suggestions, and the darker dots are the most recent ones. They suggest utilizing long photoperiods and UV periods, the success of which was confirmed in growth experiments in the Food Computer.

The most striking discovery in this experiment was the positive effect of a 24-hour photoperiod, i.e., constant daylight. This result replicated evidence on the volatile profile effects of a 24-hour photoperiod described by Skrubis et al. [39], who found that basil plants grown with a 24-hour photoperiod weighed, upon maturity, approximately 25% more than plants grown with a nine-hour photoperiod (although they took three days longer to reach maturity) and 27% more than plants grown outdoors in natural light with an approximately 15-hour photoperiod. That study also characterized changes in the relative volatile profiles of those basil plants, but not absolute volatile content, so comparisons to chemscore in the current work are not possible. The 24-hour photoperiod discovery is notable because the hand-designed experimental conditions in Round 1 had a photoperiod of 18 hours, and the experimenters and the model were blind to the Skrubis et al. study. The surrogate optimization approach nevertheless iterated the recipes into the 24-hour photoperiod, where it had a strong positive effect.

Aside from the high R-Score in Table 3, further evidence for the importance of photoperiod can be seen in the high correlation between R-Score and photoperiod in Table 4, and in the regression process itself: For each run of symbolic regression, the most parsimonious nontrivial model had the form *y = cp*, for some constant *c*, where *p* is the photoperiod. Also, Fig 4(a) shows a linear model trained on all three light variables to fit the log R-Score. Fig 4(b) shows a linear model of R-Score based on photoperiod alone. Fig 4(c) shows the predictions of a linear model trained on all three variables, but with the effect of photoperiod removed, i.e., it is trained to fit the residuals. These modeling results are similar with imputed and outlier-handled data. The low performance of the residual model suggests that photoperiod had such a dominating effect that the effects of other variables were effectively noise. However, since significant effects of UV have been reported in previous work [26,27] and are not seen here, it is also possible that there are significant nonlinear dynamics that require further trials and nonlinear modeling to uncover and exploit.

**Fig 4.**
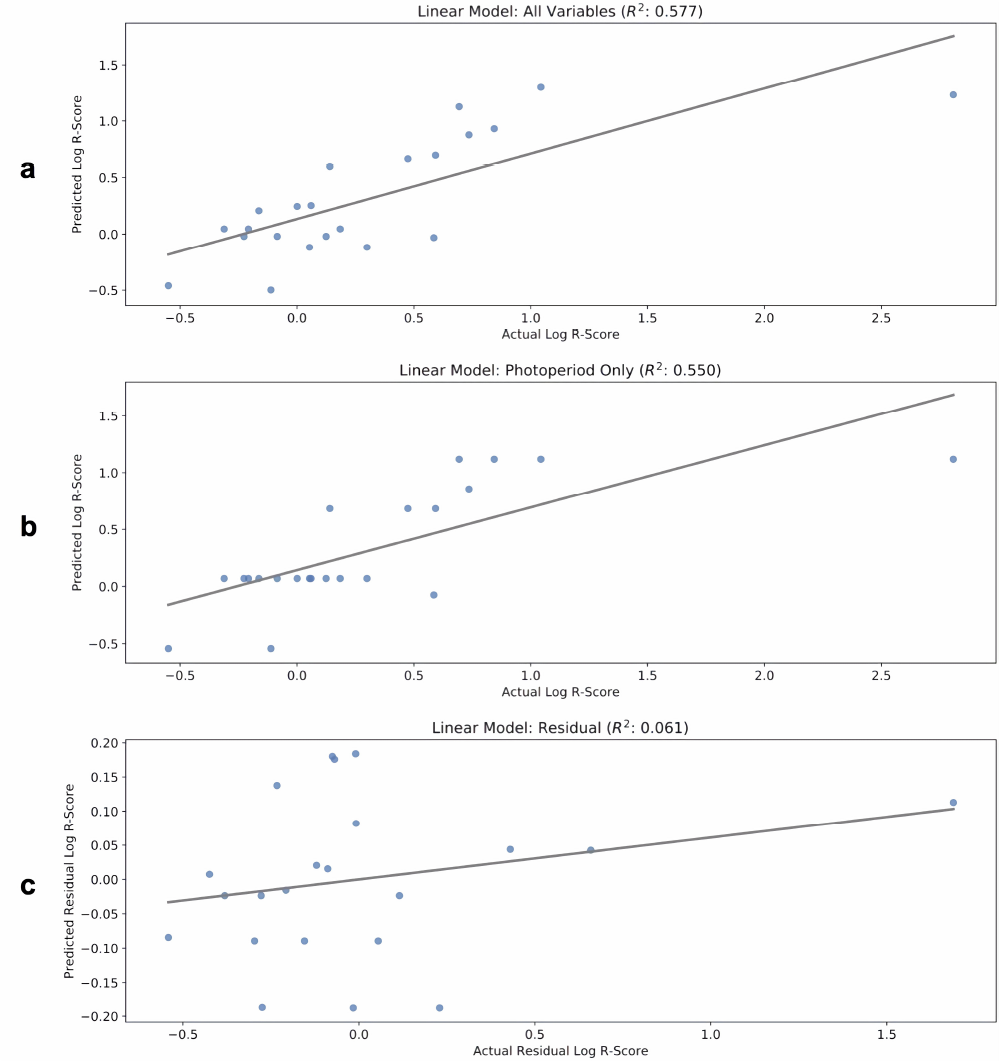
Linear regression analysis of actual vs. calculated log R-Score for three different models. (a): A linear model trained on UV, photoperiod, and PAR. (b): A linear model trained on photoperiod only. (c): A linear model trained on residuals after removing photoperiod effect. Photoperiod dominates the other variables (or possible there are significant nonlinear effects between these variables).

## 4 Discussion

The experiment described in this paper confirmed that climate recipes affect how volatile flavor molecules accumulate in basil, and that it is possible to discover good recipes iteratively through machine learning. The recipes discovered in this manner replicated known principles (such as the weight/flavor tradeoff), and also demonstrated the possibility for discovering previously unknown, surprising principles (like the 24 hr photoperiod). The 24-hour photoperiod in particular is impossible in nature (except around the summer solstice within the Arctic and Antarctic circles) and therefore unlikely to be discovered, except in controlled environments for cyber-physical agriculture.

The most immediate direction of future work is to expand the current experiment to a larger search space. A facility with four containers, making it possible to evaluate an order of magnitude more recipes at once, is in development at MIT and illustrated in Fig 5. This facility will make it possible to control a number of other actuators besides light, including temperature, pH, nutrient concentration, microbial, and other additives, and different plant cultivars. It will also be possible to measure the energy and other costs associated with the recipes, as well as objectives such as nutrient components, density, and yield, and more elements of flavor (single compounds, and ratios of compounds).

**Fig 5.**
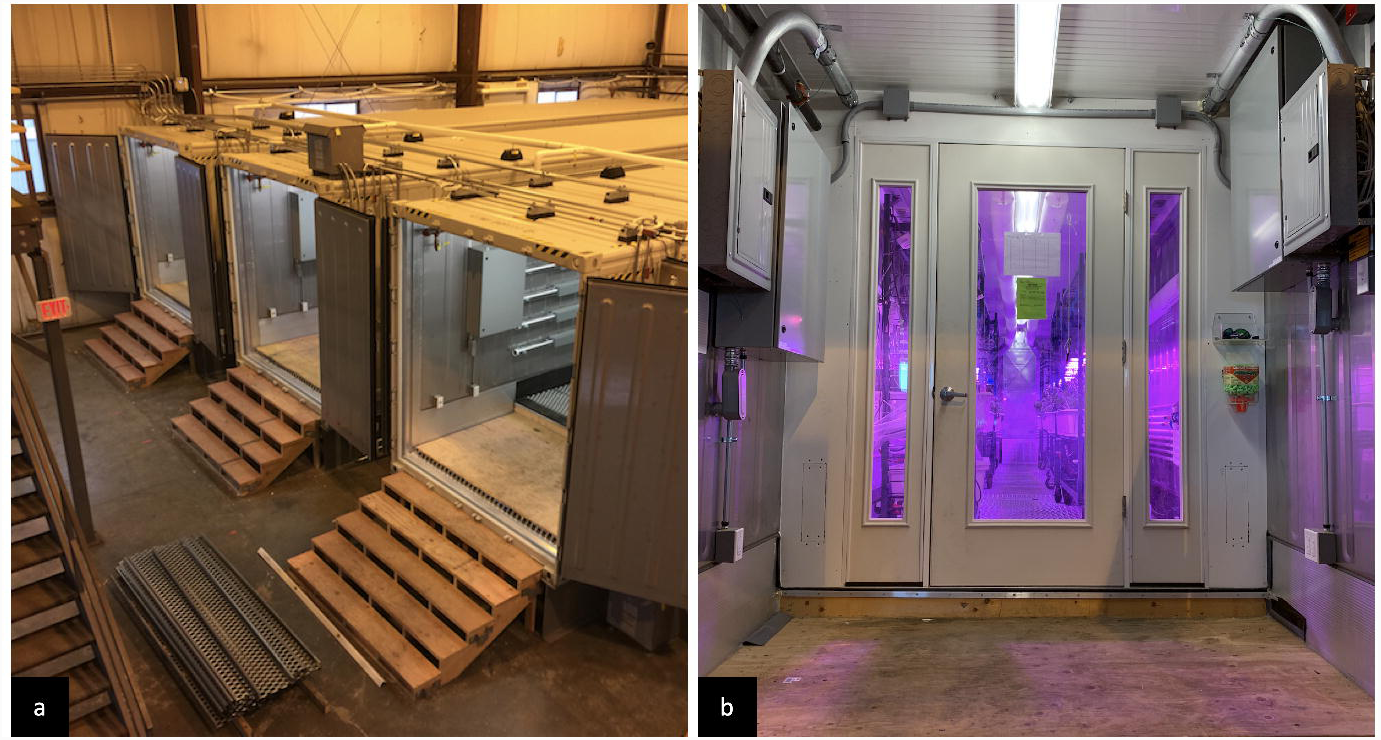
The MIT expansion facility under development. (a): Four containers being converted to large-scale Food Servers (b): The entrance to the next generation of MIT OpenAg Food Servers.

In terms of surrogate optimization, more iterations can be run to build more accurate models, and to determine the proper stopping point of the method, i.e. to run it until it has likely converged. The approach will be extended to cover the larger search space as well as multiple objectives. Most likely, different models and optimizers will be necessary. In low-dimensional settings with unknown nonlinearities and a relatively small number of samples, Kriging [34], Gaussian processes [36,49], and symbolic regression [44] are suitable choices for building a regression model of natural phenomena. When the dimensionality and number of samples increases, deep neural networks may be a better model of the solution landscape [47,50,51], and evolutionary optimization a better way to determine the most promising samples [43–45].

The next step will be to extend the experiment to other plants, such as cotton, where the goal is not to optimize flavor but physical properties such as strength and length of the fibers. It will be important to verify that such plants are viable to grow artificially, and that such properties can be optimized with available actuators, in isolation and in combination with other properties. Future extensions to other areas may include biofuels and plants with specific medicinal value.

The third future step is to extend the optimization from static recipes to time-varying recipes, i.e. optimizing the actuators during the entire growth period of the plant. Of particular interest are different stress periods when the plant is exposed to, for example, drought or signals of predators (e.g. through chitosan added to the growth medium). Such periods may produce a response in the plan that results in more flavor or more rapid growth, for example. Such recipes should be reactive, i.e. conditional to real-time measurements of the growth status. One possibility is to use machine learning to establish a mapping from visual images of the plant to more destructive measurements such as chemical concentrations. Such optimization spaces are very high-dimensional, most likely making it necessary to use evolutionary optimization, and perhaps neuroevolution to construct a mapping from sensory time series to optimal actions [55,56].

## 5 Conclusions

The experiments showed that light conditions have a large effect on the chemotype of the basil plant and that the surrogate optimization method can discover meaningful growth recipes that influence that chemotype. The results demonstrated a tradeoff between flavor and plant mass, thus confirming the well-known “dilution effect”. Furthermore, this study demonstrated how the surrogate optimization approach can discover new and unforeseen recipes that can produce better outcomes. Initially, basil was assumed to need a period of darkness in order to produce an ideal outcome, but that assumption turned out to be wrong. The highest density of flavor molecules was produced by subjecting the plants to all-day light, which the surrogate optimization approach discovered quickly and reliably. The results thus demonstrate that surrogate modeling and machine discovery can be used to find growth recipes that are both effective and surprising—and difficult and time-consuming to find through traditional hand-designed experiments.

Computer-controlled growth environments are a promising approach for the future of agriculture, potentially maximizing production and quality and minimizing waste and cost. As Food Computer technology advances, it can be useful to think of these units as a whole-plant bioreactors where experiments will contribute to the emerging field of ethnophytotechnology [57]. The initial experiments in this paper suggest that the cyber-physical approach to agriculture is indeed viable: such environments can be built, the plants thrive in them, the climate recipes make a difference in growth outcomes, and machine learning can be used to discover good recipes automatically. Future steps should verify these results on other plants, expand to larger search spaces with more actuators, and to optimizing entire growth periods. Higher-volume food computers need to be built and more powerful optimization methods employed, but the results so far suggest that such extensions are worthwhile.

## Acknowledgements

The authors would like to thank Christina Agapakis, Nate Tedford, and Scott Marr for access to and support on analysis instrumentation and Babak Hodjat and Hormoz Shahrzad for modeling and optimization insights and comments on the manuscript.

## Data availability

The data underlying the results presented in this study are freely available on the Open Agriculture Initiative’s public Github repository located at https://github.com/OpenAgInitiative/flavor-data

## Author contributions

AJJ and EM contributed to the conceptualization, analysis, methodology development, investigation, visualization, and preparing the original draft of the manuscript. AJ ran the biological and GC-MS experiments and EM ran the computational experiments. JdlP supervised and contributed to the writing, editing, interpretation of results, and review of the manuscript revision. TLS contributed to conceptualization, experimental design, fabrication of Food Computer and Food Server elements, and experimentation for biological experiments. RM and EM designed the surrogate optimization methods and EM implemented them and ran the computational experiments. CBH oversaw and contributed to conceptualization, experimental design, system design of Food Computer and Food Server elements, experimental operation, and funding acquisition.

